# Distinct nanoscale organizations of mucins and trans-sialidases in *Trypanosoma cruzi*

**DOI:** 10.64898/2026.02.23.707421

**Authors:** Gonzalo Escalante, Raquel Parada-Puig, Lucía F. Lopez, Alan M. Szalai, María de los Milagros Cámara, Luciano A. Masullo, Oscar Campetella, Juan Mucci, Fernando D. Stefani

**Author notes:** 1-Instituto de Biología y Medicina Experimental (IBYME), Consejo Nacional de Investigaciones Científicas y Técnicas (CONICET), Ciudad de Buenos Aires & Universidad de San Andrés, San Fernando, Argentina. These authors contributed equally to this work (Raquel Parada-Puig, Lucía F. Lopez, Alan M. Szalai). **Corresponding Authors:** (FDS), (JM).

## Abstract

*Trypanosoma cruzi*, the causative agent of Chagas disease, relies on surface sialylation to evade host immunity and invade cells. This process is mediated by trans-sialidases (TS) and mucins, an enzyme–substrate pair anchored to distinct lipid environments. Yet, how these molecules are organized at the nanoscale in the parasite’s membrane remains unknown. Using dual-color super-resolution microscopy, we show that ∼60% of mucins and TS are segregated into nanoclusters (∼100 nm) that rarely contact each other, distributed with a nonrandom separation distance, indicating that these abundant domains follow a specific order in the plasma membrane and are unlikely to serve as primary sites of sialylation. In contrast, the ∼40% fraction of non-clustered mucins and TS exhibits significantly shorter-than-random separation distances and appear ordered as in a shared fibrillar network. Additionally, we describe a distinct structural organization within these domains: mucins - residents of detergent-resistant domains (DRDs) - organize into high-molecular-weight complexes, whereas TS (which are excluded from DRDs) do not, suggesting a potential mechanism for domain-specific protein localization.

## Introduction

The parasite *Trypanosoma cruzi*, the etiologic agent of Chagas disease or American trypanosomiasis, presents a unique mechanism to avoid the immune response of the host that has intrigued researchers all over the world^1^. By covering its membrane with a dense coat of glycoproteins containing sialic acids, the parasite avoids lysis by serum factors^2,3^ and achieves the invasion of the host cell^4^. Remarkably, *T. cruzi* is unable to synthesize sialic acids. Instead, *T. cruzi* is part of a restricted group of parasites evolutionarily adapted to scavenge sialic acid from exogenous sialoglycoconjugates by means of a modified α(2,3)-sialidase known as trans*-*sialidase (TS), a glycosylphosphatidylinositol (GPI)-anchored protein present on the cell surface of trypomastigotes^5^ and actively released from the parasite’s plasma membrane in microvesicles^6^. In the presence of glycans with an α(2–3)-sialyl residue linked to a terminal βgalactose, the parasite’s surface becomes rapidly sialylated through a glycosyl-transfer reaction driven by TS^7^. Although probably not the unique acceptors available, mucins are considered the main targets for TS-transferred sialyl residues^2,8,9^. Mucins, with typical sizes ranging from 60 to 200 kDa depending on the parasite strain, are a group of surface proteins, also GPI-anchored, that feature multiple *O*-linked oligosaccharide chains and are highly immunogenic^10^ .

The genome of *T. cruzi* encodes approximately 1,400 gene members of the TS-like superfamily^11^ and 500 to 700 genes of the mucin family^12^. This diversity is key for the parasite’s virulence by enhancing its capacity to modify its surface and evade host immune defenses^13,14^. Mucins and TS present well-distinct tertiary structures: the core of TS is the typical of a globular glycoprotein, while mucins are highly *O*-glycosylated stick proteins^14^.

Prior research revealed that TS and sialylated mucins, despite both being GPI-anchored, appear in separated domains of the membrane of *T. cruzi* trypomastigotes^6,15^. The discovery of the segregated distribution of mucins and TS raised significant questions regarding the mechanisms that regulate their compartmentalization and interaction^6^. However, the insights provided by these early studies were limited because the spatial resolution achieved did not allow for detailed characterization of the size or spatial organization of mucins and TS domains.

It has also been established that mucins and TS membrane domains exhibit distinct biochemical organizations, with mucin domains possessing lipid raft characteristics, while TS domains lack them^6,15^. Notably, this correlates with the fact that, in the mammalian stage of the blood trypomastigote, these proteins associate with different lipids: mucins are bound to an alkyl-acylglycerol-containing GPI, whereas TS bear ceramide^16,17^. While in mammalian cells ceramide has not been found in GPI-anchored proteins, alkyl-acylglycerol (and diacylglycerol to a lesser degree) is a key determinant of their association with lipid rafts^18,19^. Therefore, this distinct lipid composition could be a possible explanation for the differential membrane organization.

Here, we apply two-color super-resolution fluorescence microscopy combined with clustering analysis, molecular distance quantification, model-based simulations and biochemical approaches to resolve the nanoscale landscape of TS and sialylated mucins on the surface of *T. cruzi* trypomastigotes with unprecedented detail. This integrated approach allows us to test whether the long-standing view of mucin–TS interactions—largely inferred from biochemical assays and diffraction-limited imaging—holds true at the nanometer scale between this enzyme-substrate pair. By directly quantifying cluster sizes, spatial distributions, molecular proximities, and the underlying structural dimensionality, we uncover architectural principles that reshape our understanding of how these key virulence factors are organized and potentially coordinated on the parasite membrane. Furthermore, Blue Native non-denaturing Polyacrylamide Gel Electrophoresis (BN-PAGE) analysis reveal distinct structural organization within mucins and TS domains, were DRDs-resident mucins appear to assemble in high-molecular-weight complexes, while non-DRDs-resident TS do not. Ultimately, this framework enables us to propose spatially grounded hypotheses for protein distribution on the plasma membrane and suggest how sialylation can occur *in vivo*, providing important mechanistic insights into host–parasite interactions.

## Results

### Nanoscale distribution of mucins and trans-sialidases domains

We applied two-color Stochastic Optical Reconstruction Microscopy (STORM)^20,21^ to fixed *T. cruzi* cells in the trypomastigote stage. The SAPA region of TS—*C*-terminal domain composed essentially of immunodominant amino acid repeat units^22^—was immunoconjugated to Alexa Fluor 568. Sialylated mucins were labeled with Alexa Fluor 647. To label sialylated mucins, we followed an unnatural sugar approach that enables labeling TS target proteins with higher selectivity and minimal background compared to the use of mucin-directed antibodies or Lectins^7^.

Super-resolution imaging revealed that an important fraction of mucins and TS are organized in separated domains of nanometric dimensions (Figure 1A). By contrast, conventional diffraction-limited microscopy only provides rough signs of segregation for these two proteins (Figure 1B), as previously reported^6^.

**Figure 1.**
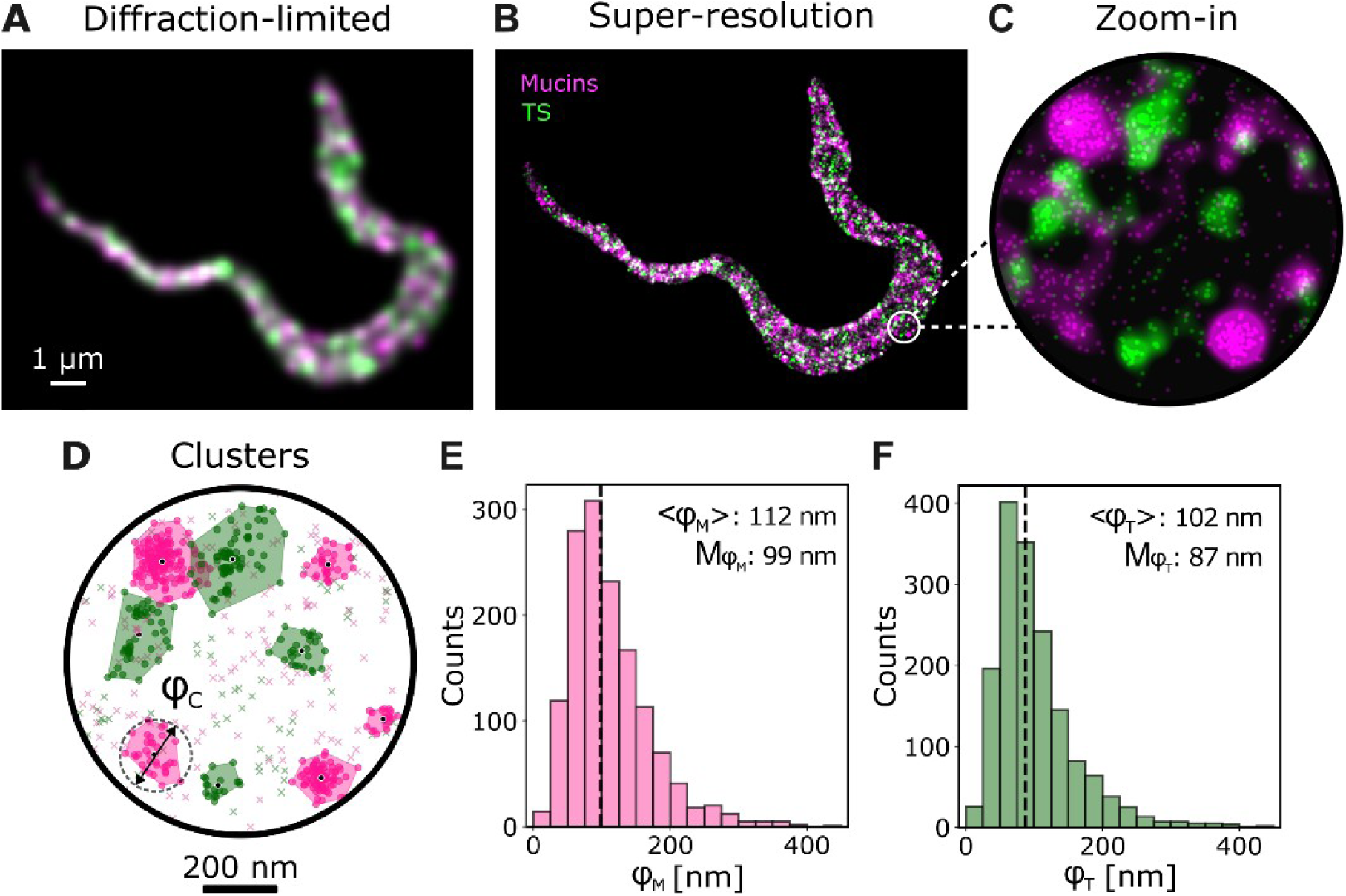
Mucins and TS in T. cruzi plasma membrane. (A,B) Fluorescence imaging of mucins (magenta) and TS (green) in a *T. cruzi* trypomastigote – (A) diffraction-limited wide-field image, (B) Two-color super-resolution image based on single molecule localizations (STORM). The image is rendered by plotting each localization as a two-dimensional gaussian function with standard deviation of 10 nm. (C) Zoom-in of the super-resolved image of mucins and TS on the membrane region marked in (B) overlayed with the individual molecular localizations. (D) Example of cluster analysis using DBSCAN in the same region shown in (C). Localizations classified as clustered are represented with dots while non-clustered localizations are represented with crosses. The estimation of the effective diameter of the clusters (φ_C_) is depicted. (E, F) Distributions of effective cluster diameters of mucins (E, φ_M_) and TS (F, φ_T_). The average effective diameters are indicated in brackets, while the medians are marked with dotted lines and represented as Mφ.

To analyze the spatial distribution of the molecular localizations, we employed a density-based clustering algorithm (DBSCAN)^23^ and a density-based clustering validation^24^ to detect clusters of mucins and TS. Overall, 184 membrane areas were analyzed from 25 trypomastigotes. 1,617 clusters of TS and 1,413 clusters of sialylated mucins were identified, with heterogeneous sizes and shapes. The cluster size was quantified as the diameter of the smallest circle including the cluster (Figure 1C). The histograms of cluster size reveal monomodal, positively skewed distributions with mean values of 110 and 100 nm for mucins and TS, respectively (Figures 1D and 1E). The median cluster diameter for mucins and TS were determined to be 100 nm and 90 nm, respectively, in agreement with previous results using single color GSDIM super-resolution microscopy^6^. These sizes are well below the diffraction limit and are comparable to the sizes of other reported clusters of membrane proteins such as T-cell receptors (from 70 nm to 140 nm)^25^ and to the reported sizes of lipid rafts (up to 200 nm)^26^.

Several studies have established that crosslinking agents can potentially generate or stabilize protein clusters^27–30^. In our previous work, where the membrane segregation of mucins and TS was first detected^6^, numerous fixation reagents and temperatures were tested in order to confirm the biological origin of the observed protein organization. Moreover, a rapid membrane fluidization using diethyl-ether abolished the distinct pattern observed^6^. By detecting a uniform distribution of the highly abundant Variant Surface Glycoprotein (VSG) in *Trypanosoma brucei*—using the same fixation protocol as in our measurements of mucins and TS in *T. cruzi* tissue-culture-derived trypomastigotes—we further confirm that the observed nanodomains are not fixation artifacts (Supplementary Figure 1). Moreover, we also note that the maximal size reported of fixation-induced protein clusters after *p*-formaldehyde (PFA), glutaraldehyde (GA), or methanol (MeOH) is of 14–16 nm in diameter^29^. In contrast, the domains revealed in this work present mean effective diameters about an order of magnitude larger. All the aforementioned supports the idea that fixation is not responsible for the observed spatial organization of mucins and TS.

Once these nanodomains were identified, a relevant question arises as to whether they are randomly distributed or if they follow any other underlying organization. To address this question, we calculated the distribution of first neighbor distances between the centers of mass of the experimentally observed clusters of mucins (d_MM_) and TS (d_TT_), and compared those distributions to the ones obtained from the same clusters in randomized positions within the same observation areas (Figure 2A). 100 random distributions were simulated for each one of the experimentally observed sets of clusters. The distributions and cumulative distribution functions (CDF) of d_MM_ and d_TT_ are shown in Figures 2B and 2C, respectively, for the experimentally observed and the randomized distributions of clusters. For the randomized distributions, all the individual CDFs are shown (grey) along with their average (black). Remarkably, neither mucins nor TS clusters follow a random distribution over the parasite membrane. The first neighbor distances between mucin clusters are larger than those of a random distribution, indicating that these clusters are segregated. By contrast, TS clusters are found at shorter distances compared to those expected in a random distribution. These observations, supported by a two-sample Kolmogorov-Smirnov test (p-value: 1.4×10^-3^ for d_MM_; p-value: 1.1×10^-9^ for d_TT_), indicate that the clusters of mucins and TS are not randomly distributed but instead reflect an underlying membrane organization.

**Figure 2.**
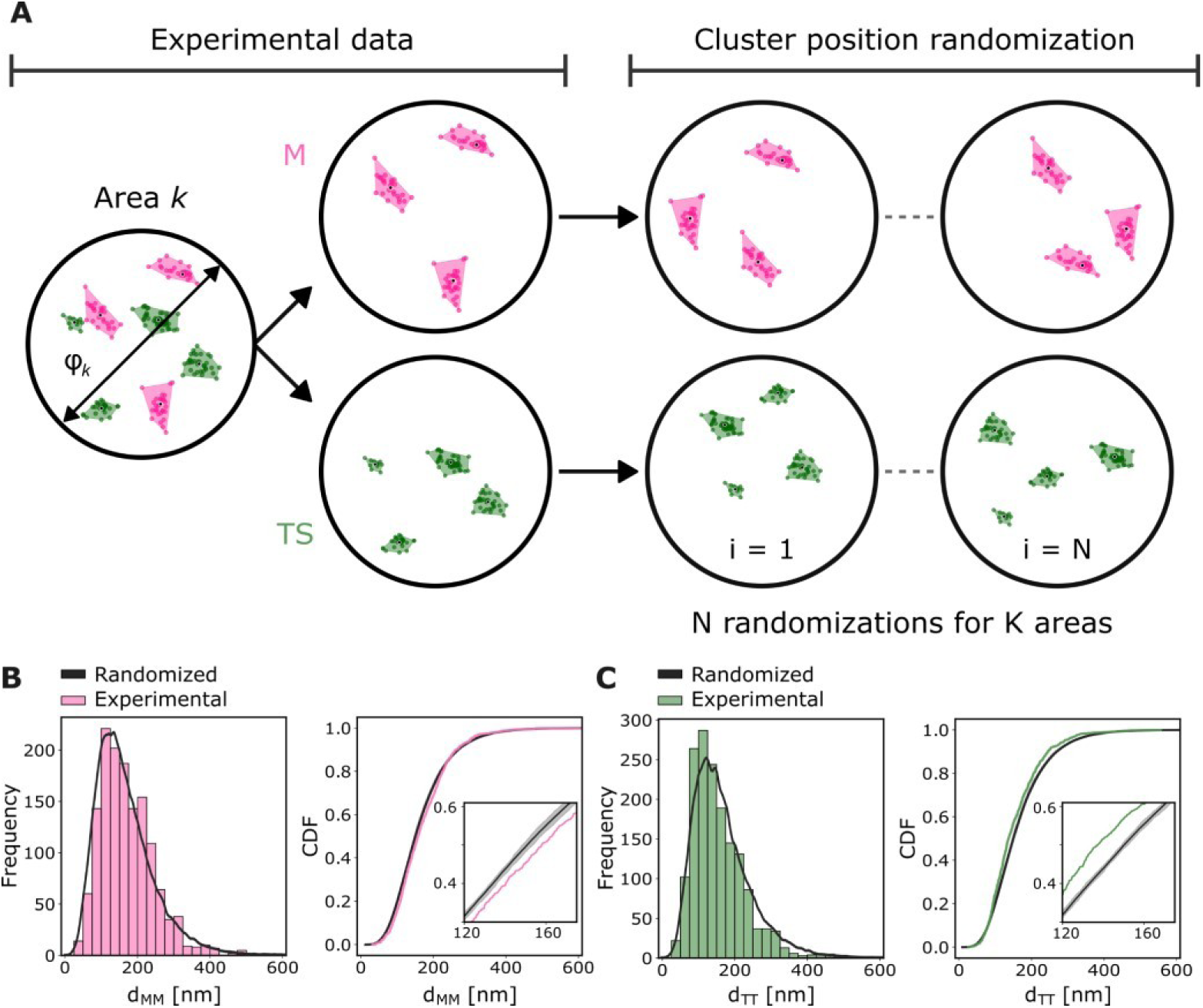
Characterization of the spatial distribution of mucins and TS nanodomains. (A) Schematic description of the cluster position randomization: within each experimentally observed area, the clusters found with DBSCAN were relocated in random positions ensuring that they do not overlap to avoid cluster merging. (B, C) Histogram and Cumulative Distribution Function (CDF) of first neighbor distances between the mass centers of experimental and randomized clusters of mucins (B, d_MM_) and TS (C, d_TT_). The CDFs of 100 randomized distributions are displayed in grey, with their average highlighted in black. A zoom-in is shown in the insets to better appreciate the differences between the curves.

### Separation distances and overlap between mucins and TS domains

Next, we analyzed the nearest-neighbor cross distances and overlap between clusters of mucins and TS. Because both populations exhibit intrinsic spatial organization and are not randomly distributed (see previous section), a conventional comparison with fully random cluster distributions would not provide an appropriate reference. Instead, to generate a realistic null model in which mucin and TS clusters are spatially independent while preserving the native organization of each population, we implemented a shuffling procedure. Specifically, experimentally determined mucin clusters from one region were paired with TS clusters from a different region. This procedure preserves the intrinsic spatial organization of each cluster population while removing any local spatial correlation between them (Figure 3A).

**Figure 3.**
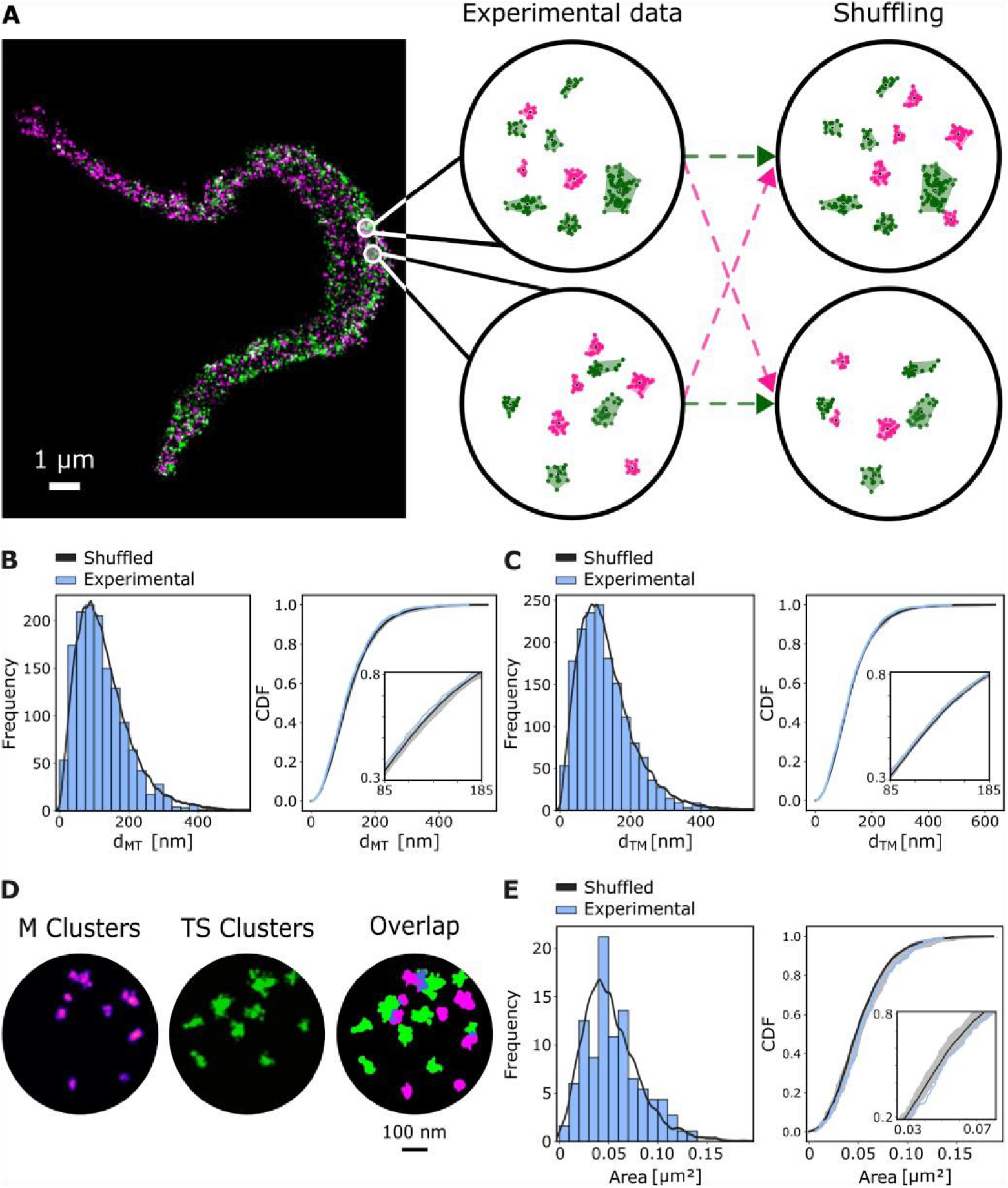
Separation distances and overlap between mucins and TS nanodomains. (A) Shuffling of experimental data: given two circular areas of equal diameter of *T. cruzi* membrane, mucin clusters from one area are combined with TS clusters from the other (and vice-versa). First neighbor distances from the mass centers of mucins clusters to TS clusters (d_MT_) and vice versa (d_TM_) are determined. (B, C) Histogram and CDF of first neighbor cross-distances (B, d_MT_ –C, d_TM_). For each case, the CDFs of 100 randomized distributions are displayed in grey, with their average highlighted in black. (D) Example of 2D Gaussian rendering of an experimental cluster dataset: mucins clusters in magenta, TS clusters in green, overlap areas in blue. (E). Histogram and CDF of overlap areas between mucins and TS clusters for experimental and shuffled data.

Using this approach, we generated the distributions expected under the null hypothesis that mucin and TS clusters are independently organized. The experimentally observed nearest-neighbor distance distributions from mucin clusters to TS clusters (p = 0.06) and from TS clusters to mucin clusters (p = 0.33) were not significantly different from those obtained from the shuffled datasets (Figures 3B and 3C). These results indicate that the measured inter-cluster distances can be explained by the intrinsic spatial organization of mucin and TS clusters and do not provide evidence for additional spatial coupling between the two populations.

To evaluate the spatial overlap between mucin and TS domains, localization maps were first rendered as two-dimensional Gaussian images and subsequently converted into binary masks. Overlap areas were quantified by calculating the number of pixels common to the mucin and TS masks (Figure 3D). Consistent with the nearest-neighbor analysis, the experimentally observed overlap distribution did not differ significantly from that obtained using the shuffled null model (Figure 3E; K-S statistic = 0.09, p = 0.09), indicating that the degree of overlap is also consistent with the independent spatial organization of the two cluster populations.

### Non-clustered mucins and TS arrangement

The spatial analysis also reveals that 38% of the mucins and 42% of the TS are located outside clusters. The distributions of first neighbor distances between non-clustered mucins (M-M) or non-clustered TS (T-T) do not provide any insight into their molecular distributions because they both present a peak at the average single molecule localization precision (∼10 nm) (Figure 4A), which is explained by multiple localizations of the same fluorophore (multiblinking)^31^ and/or multiple labeling of the same protein. On the other hand, the cross-distance distributions, from mucins to TS (M-T) and *vice versa* (T-M), are of interest and present broader distributions and with larger mean values of 35 nm and 45 nm, respectively (Figure 4B).

**Figure 4.**
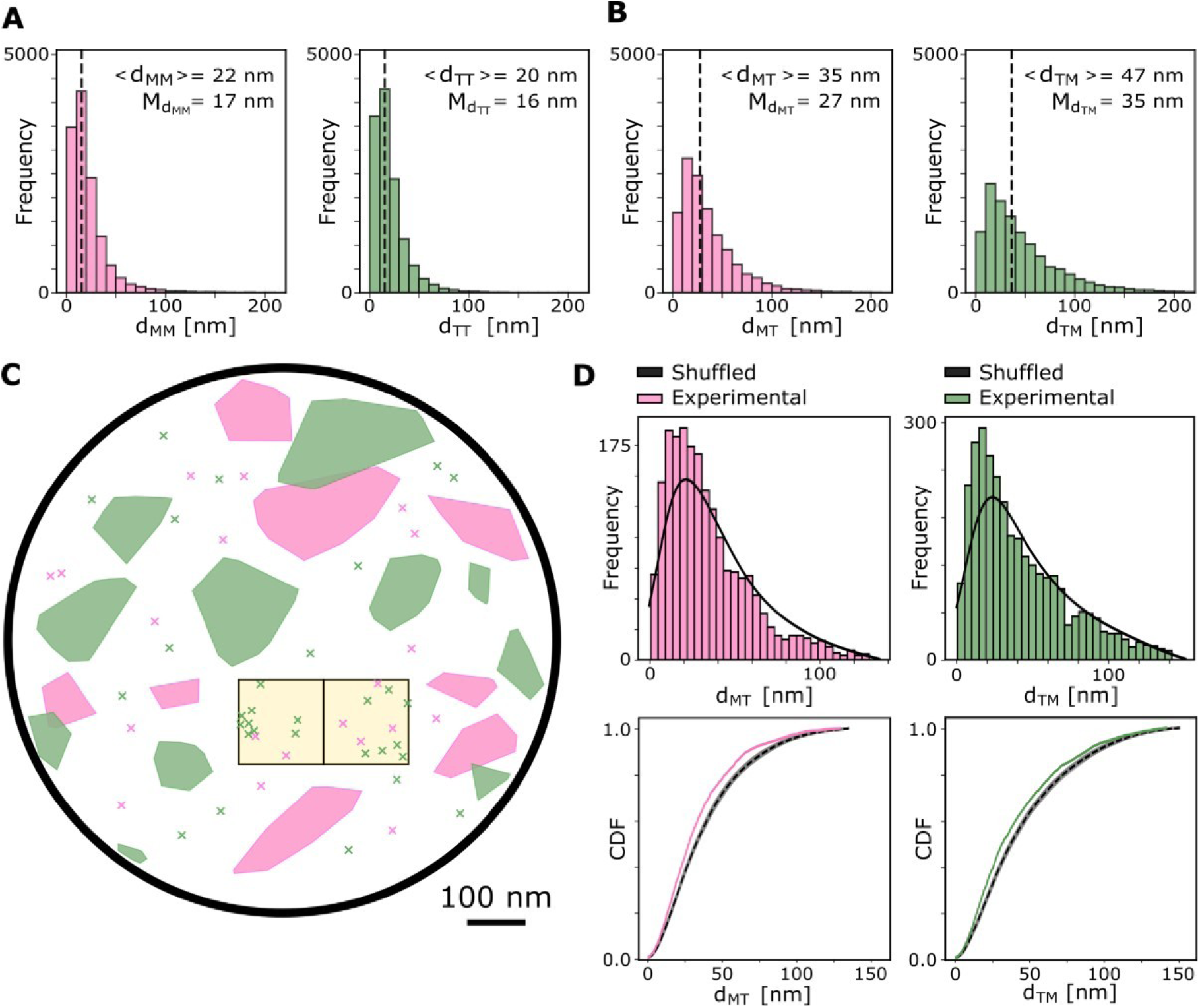
Analysis of the spatial distribution of non-clustered mucins and TS molecules. (A,B) Histograms of the first neighbor distances of non-clustered proteins of the same kind (A, d_MM_ and d_TT_) and between mucins and TS (B, d_MT_ and d_TM_). Mean values are reported in brackets and the median is shown as a dashed line. (C) Schematic of the sampling of the membrane left available by the clusters using square areas. In each one of the 184 circular areas analyzed, the membrane area not occupied by clusters was sampled by taking non-overlapping, non-repetitive squares with a side of 146 nm, ensuring that they contained both mucins and TS localizations. (D) Histogram and CDF of experimental d_MT_ (magenta) and d_TM_ (green) compared to the distance distributions obtained from shuffling mucins and TS localizations from different square areas (in black). Kolmogorov–Smirnov test p-values are 2.5×10^-15^ for M-TS and 3.9×10^-14^ for TS-M.

The question then arises of whether these distributions are simply a consequence of the overall abundance of each protein and random placement over the available space, or if they reflect some other underlying organization. We thus compared the experimental distributions to simulated random distributions, considering the same number of experimental localizations and the available space left by the clusters. In order to maintain the original distributions of non-clustered localizations of mucins and TS, we applied the same shuffling method used for the clusters. To implement this approach, we sampled the membrane not occupied by clusters with square regions, as shown in Figure 4C, ensuring that each square area contained both mucins and TS molecules. The size of the square region was chosen as the minimum that encloses the maximum first neighbor distances observed experimentally for M-T and T-M (∼200 nm, Figure 4B), namely 146 nm x 146 nm squares with a diagonal of 206 nm. In this way, we could maximize the area of non-clustered proteins analyzed while respecting the maximum first neighbor distances observed experimentally.

Figure 4D shows the distributions and CDFs of first neighbor cross distances for the native distribution of non-clustered mucins and TS molecules, and for the randomized (shuffled) localizations obtained from combining mucins and TS from different areas. In both cases (M-T and T-M distances), the shuffled localizations present overall larger distances, indicating that the non-clustered mucins and TS do not occupy the entire available membrane. Instead, they are confined to smaller compartments or structures that shorten the separation distances to their nearest counterpart.

A way of gaining insight into the spatial confinement of the non-clustered mucins and TS is by studying the dimensionality (*n*) of the underlying space in which these localizations reside^32^. This approach is based on examining the distance dependence of the complementary cumulative distribution function (CCDF) of the nearest-neighbor distance, and enables determining the basic organization of molecules even in highly sparse conditions^32^. Molecules randomly organized in one dimension (1D) present a value of *n* = 1. Molecules distributed randomly in 2D yield a value of *n* = 2. Distributions where molecules are organized in confined structures (e.g., fibers of finite width) can attain values of *n* between 1 and 2 (Figure 5A)^32^. Performing this analysis to the localizations of non-clustered mucins and TS reveals that both proteins show the same average value of *n =* 1.36 (Figure 5B), confirming the confinement and suggesting that they may be sharing a common compartmentalization.

**Figure 5.**
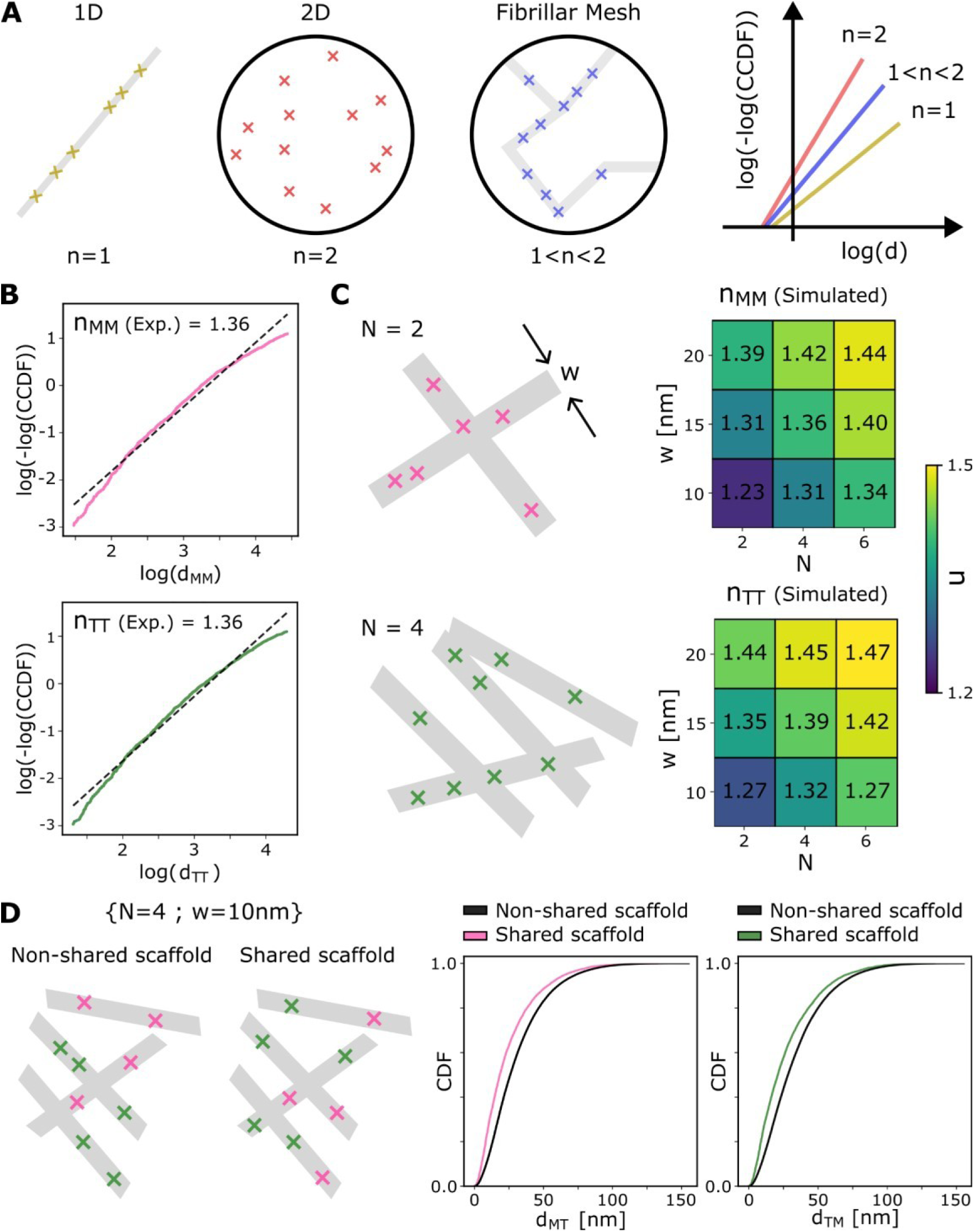
Dimensional analysis of non-clustered mucins and TS molecules. (A) Functional dependence of the complementary cumulative distribution function (CCDF) of the distance to the first neighbor (d) for various point distributions. The log[-log(CCDF)] as a function of the log(d) is a linear function whose slope *n* (dimensionality) takes extreme values of 1 for a strictly linear arrangement (1D) and 2 for a random 2D distribution. Intermediate organizations such as fibrillar networks present values of 1 < n < 2. (B) log[-log(CCDF)] vs. the experimentally observed first neighbor distances of mucins (d_MM_) and TS (d_TT_). Dotted lines are linear fits retrieving a value of *n* = 1.36 in both cases. (C) Schematic of molecules distributed over a mesh of N rectilinear fibers of width (w). Average values of n obtained from simulations randomly distributing a density of molecules equal to the experimentally observed for mucins and TS over N (2, 4, or 6) rectilinear fibers of various thicknesses (10, 15 or 20 nm) inside an area of 146 nm x 146 nm. For each condition, 10,000 simulations were performed. Results for other combinations of N and thickness are presented in Supplementary Figures 2B and 2C. (D) Schematic representations illustrate two scenarios: mucins and TS arranged on separate fibrillar scaffolds versus both sharing a common scaffold. The plots show cumulative distribution functions (CDFs) of first-neighbor cross-distances derived from simulations. In these simulations, molecules were randomly distributed at densities matching experimental observations across a mesh of four rectilinear fibers (10 nm wide) within a 146 × 146 nm area. Magenta and green curves represent CDFs of distances from mucins to TS and TS to mucins, respectively, when both proteins share the same scaffold. The black curve corresponds to the scenario where mucins and TS occupy distinct fibrillar scaffolds.

Using simulations, we tested three models of compartmentalization of the non-clustered mucins and TS: circular compartments, rectilinear fibers, and curved fibers. The simulations consisted in: (i) defining compartments within square regions with a side of 146 nm (as the ones experimentally sampled), (ii) placing mucins and TS at random positions within the compartments so as to obtain an overall surface density equivalent to the experimentally observed, and (iii) computing *n*. In the case of the circular compartments the position, number and diameter of the compartments were varied. In the case of fibers, the number (N), position, orientation and width (w) of the fibers were varied (Figure 5C). Interestingly, it was not possible to reproduce the experimentally observed dimensionality of *n* = 1.36 with the model of circular compartments, regardless of the compartment parameters used (Supplementary Figure 2A); this model led to values of n closer to the extreme values of 1 or 2, and was therefore ruled out. By contrast, both fibrillar models retrieved values of dimensionality comparable to the experiments (*n =* 1.30 - 1.42) for several combinations of 2 < N < 6 and 10 < w < 20. Figure 5D shows the results for the model of rectilinear fibers (see Supplementary Figures 2B and 2C for further results).

A more direct comparison to the experiments can be obtained by evaluating the CDFs of first neighbor cross distances of simulations where molecules were randomly distributed over a mesh of 4 rectilinear fibers of 10 nm width inside an area of 146 nm x 146 nm, while keeping the overall density of molecules equal the experimentally observed (further results for the curved fibers and linear fibers in Supplementary Figures 3 to 6). In this way, we generated variations of molecular distributions with a similar density and dimensionality (n) as the experimentally observed for mucins or TS. Then, we computed the CDF of first neighbor cross distances in two scenarios: (i) mucins and TS distributed over the same fibers and (ii) mucins and TS distributed on different fibrillar scaffolds (Figure 5D). The latter was achieved by shuffling mucin and TS positions from different simulations, just as it was done with the experimental data from different regions. The cross-distances to the nearest neighbor increased when mucins and TS were distributed in independent fibrillar meshes, just as it was observed when the experimental data was shuffled for randomization. This result strongly suggests that the positions of both proteins are determined by the same underlying structure. Otherwise, if both proteins were already distributed in independent structures with a dimensionality of *n* = 1.36, shuffling their relative positions should not have an effect on their relative separation.

### Native protein oligomerization underlies differential membrane partitioning

Disentangling the biomolecular mechanisms and functions underlying the peculiar nanoscale organization of mucins and TS on the membrane of *T. cruzi* will require substantial further investigation. As an initial step in this direction, we examined the native oligomerization states of mucins and TS, given that protein oligomerization is a well-established regulatory mechanism controlling the sorting, membrane partitioning, and spatial stabilization of GPI-anchored proteins in mammalian cells.^33–35^ This raises the possibility that similar principles may contribute to the organization of surface molecules in *T. cruzi*.

Native protein assemblies were examined by Blue Native Polyacrylamide Gel Electrophoresis (BN-PAGE) followed by Western blotting. Tissue-culture-derived *T. cruzi* trypomastigotes were lysed under mild detergent conditions, designed to preserve protein–protein interactions. As a control for native oligomeric resolution, we included variant surface glycoprotein (VSG) from *Trypanosoma brucei* bloodstream-forms, a well-characterized dimeric GPI-anchored protein with molecular weight of ∼100 kDa^36^. Detection of VSG dimers confirms the proper electrophoretic run and the non-denaturing separation of proteins (Figure 6A).

**Figure 6.**
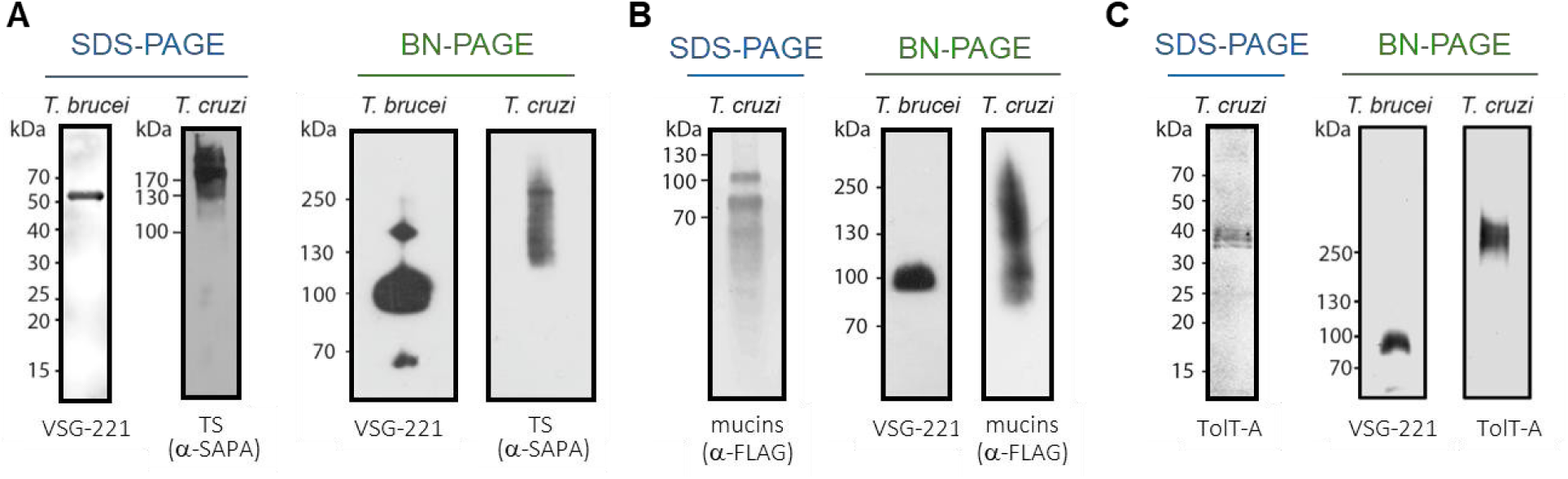
Comparative analysis of the native structural organization of membrane protein domains. Migration patterns between (A) TS (α-SAPA), (B) labeled mucins (α-FLAG) and (C) TolT-A (α-TolT-A) proteins were compared by Western blot (WB) under non-denaturing conditions (BN-PAGE, 3-12% gradient). *T. brucei* VSG glycoprotein (α-VSG) was included in each run as dimeric control (∼100 kDa). For each lane, proteins corresponding to 10^7^ tissue-culture-derived *T. cruzi* trypomastigotes (WT, CL Brener strain) or 1 x 10^6^ bloodstream *T. brucei* trypomastigotes (427 strain) were used. SDSPAGE runs are included in each case for comparison under denaturing conditions (left panels in each case).

Under non-denaturing conditions, TS displayed migration profiles comparable to those observed in denaturing SDS-PAGE, with no detectable signal at high molecular weight (Figure 6A). This behavior indicates that TS does not form stable higher-order oligomeric assemblies in the parasite membrane. In contrast, sialylated mucins exhibited prominent high-molecular-weight species in BN-PAGE (∼100 to 250 kDa - Figure 6B), migrating well above their apparent molecular weight in SDS-PAGE (70 to 100 kDa).^6,7,37^ Following the same experimental approach, we also analyzed the oligomerization state of TolT-A protein, another GPI-anchored membrane glycoprotein^38^ and DRD-resident^6^. A pattern similar to mucins was observed: under native conditions TolT-A migrated predominantly as high-molecular-weight assemblies (>250 kDa), despite its relatively low molecular weight (35-40 kDa) under denaturing conditions (Figure 6C).

Together, these data reveal a marked difference in the native supramolecular organization of membrane proteins localized to distinct lipid environments. GPI-anchored proteins associated with detergent-resistant domains exhibit stable higher-order assemblies, whereas the non-detergent-resistant resident TS does not display evidence of oligomerization under the conditions tested.

## Discussion

Our results provide new insights into the nanoscale spatial organization of mucins and trans-sialidases (TS) in trypomastigote membranes. Using two-color super-resolution microscopy and spatial analysis of single-molecule localizations, we identified that mucins and TS are present in at least two distinct spatial states. First, consistent with previous reports^6^, we observed that both mucins and TS form nanoclusters in separate membrane regions. The higher resolution achieved in our study allowed us to resolve their sub-diffraction-limited sizes, around 100 nm, which is consistent with the dimensions of other reported protein clusters and lipid rafts. Second, we found that approximately 40% of the mucins and TS molecules are distributed diffusely across the membrane, outside of clusters.

Mucins and TS share several key features. Both proteins are GPI-anchored and reach the plasma membrane via common trafficking pathways. Also, the observed clusters of mucins and TS are similar in size. However, the spatial organizations of mucin and TS clusters differ markedly. In comparison to a random distribution, mucin clusters are more widely spaced, while TS clusters are more closely packed. These findings indicate that these two proteins interact with an additional layer of heterogeneity in the plasma membrane, consistent with their differential distribution across distinct lipidic domains. Using BN-PAGE, we show that mucins and TolT-A (both DRDs-residents) assemble into high-molecular-weight complexes under non-denaturing conditions, whereas TS (non-DRD) remains largely monomeric. This structural asymmetry provides a plausible mechanistic pathway for their differential nanoscale organization in clusters observed by super-resolution microscopy. More broadly, these observations support a model in which membrane heterogeneity arises from the interplay between lipid environment and protein supramolecular assembly. Similar principles have been described for GPI-anchored proteins in polarized mammalian cells^33–35^, suggesting that oligomerization-dependent partitioning may represent a conserved strategy for organizing cell surfaces across eukaryotes.

Previous studies proposed that the spatial separation of mucin and TS clusters could hinder efficient sialylation^14^ and that the shed enzyme could be the main source of trans-sialidase activity^6^. Our results support this interpretation. Given that the catalytic reach of TS is ∼50 nm, we computed that only ∼15% of mucin–TS cluster pairs fall within a range that could theoretically enable sialylation—and even then, only peripheral molecules would be accessible. Furthermore, the observed overlap between mucin and TS clusters is consistent with random contacts, thus supporting the hypothesis that clustered states are unlikely functional sites of sialylation.

A different situation arises from the spatial organization of the ∼40% of mucins and TS that are non-clustered over the membrane, which exhibit (i) mucin-TS separation distances closer than random, and (ii) a distribution consistent with both proteins located over a common fibrillar mesh. The nearest-neighbor distance distributions from non-clustered mucins to TS and from TS to mucins exhibit median values of 35 nm and 47 nm, respectively. Although more than 60% of these non-clustered molecules lie within the theoretical catalytic range for sialylation, directly testing the occurrence of sialylation at the membrane within these regions remains challenging, given that the substantial amount of TS shed by the parasite could mask the phenomenon. Consequently, this interpretation provides the groundwork for a plausible hypothesis to be examined in subsequent studies, rather than an absolute conclusion.

Our work provides the first nanoscale map of mucins and TS on the *T. cruzi* membrane, revealing two coexisting domain organizational modes. Approximately 60% of both proteins form segregated nanoclusters with distinct (i) non-random distributions and (ii) spatial separations that make catalytically meaningful interactions highly unlikely. Meanwhile, the remaining ∼40% of non-clustered mucins and TS display a common spatial organization with the majority of molecules within the theoretical catalytic radius for sialylation. Moreover, we show that the oligomerization state plays a significant role in the organization of protein domains, with proteins localized to DRDs exhibiting higher oligomerization states—a characteristic that has been consistently associated with DRD partitioning in mammalian cells.

By linking native supramolecular organization to nanoscale membrane architecture, our findings provide mechanistic insight into how the *T. cruzi* surface may accommodate both segregation and functional interaction between key virulence factors. The revealed organizational logic of mucins and TS likely contributes to the regulation of host–parasite interactions, highlights general principles through which complex membrane landscapes are established, and enables the postulation of novel hypotheses. For instance, the clusters may fulfil functions related to the interaction/adhesion of the parasite to the host cell, to the secretion of mucins and TS to the extracellular medium, or act as reservoirs of mucins and TS that dynamically leave the clusters to bind to a common fibrillar subjacent structure. The nonclustered mucins and TS could simply be proteins en route to incorporation into functional clusters, or participate actively in the catalytic sialylation. Future studies may focus on testing these hypotheses. Methodologically, this work highlights the power of combining nanoscale imaging, spatial analysis and modelling with biochemical approaches to uncover organizational principles that are otherwise invisible, yet central to the biology of protozoan pathogens.

## Materials and Methods

### Ethics statement

The protocol of animal immunization followed in this study was approved by the Committee on the Ethics of Animal Experiments of the Universidad Nacional de San Martín, according to the recommendations of the Guide for the Care and Use of Laboratory Animals of the National Institutes of Health.

### Parasites cultures

*T. cruzi* CL Brenner tissue-culture-derived trypomastigotes were obtained from infected Vero cells growing at 37°C and 5% CO_2_ in minimum essential medium (MEM) supplemented with 4% (v/v) fetal bovine serum (FBS), 0.292 g/l L-glutamine, 100 IU/ml Penicillin and 100 μg/ml Streptomycin (all from Gibco, NY).

### Sialic acid acceptor molecules labeling

Sialic acid acceptors on the surface of *T. cruzi* trypomastigotes were labeled via the Staudinger-Bertozzi click-chemistry approach, employing the trypomastigote endogenous TS^6^. Briefly, tissue-culture-derived trypomastigotes were harvested by centrifugation, washed twice with Phosphate Buffered Saline (PBS, 137 mM NaCl, 2.7 mM KCl, 1.8 mM KH_2_PO_4_, 10mM Na_2_HPO_4_, pH 7.4; Gibco) and resuspended at a density of 1 x 10^6^ cells/μl. Parasites were then incubated in PBS with 1mM Neu5Az-GalβOMe (Neu5AzGal was synthesized following previously established methods^39^ and then washed with PBS. Afterwards, sialylated parasites were incubated with 50 μg/ml of FLAG conjugated dibenzocyclooctyne (DBCO(PEG)_4_-FLAG, Jena Bioscience, Germany) for the click-chemistry reaction. Following two additional PBS washes, labeled parasites were fixed with 4% PFA in PBS. All three incubations were 30 min long and conducted at room temperature.

### Immunostaining

Tissue-culture-derived trypomastigotes—labeled as described above—as well as exponentially grown epimastigotes and *T. brucei* bloodstream trypomastigotes were collected by centrifugation and extensively washed with PBS. Parasites were then resuspended in PBS containing 4% (v/v) *p*-formaldehyde (PFA, Electron Microscopy Sciences, PA) and fixed for 30 min, followed by washing and resuspension in PBS. Fixed parasites were settled on polyL-lysine-coated coverslips (16 or 22 mm Φ; Marienfeld, Germany) for 1 h. Unattached cells were then removed by washing with PBS. Subsequently, samples were incubated 30 min in a blocking solution (BS) consisting of 0.45 μm-filtered 5% (w/v) biotechnology grade-bovine serum albumin (BSA, Amresco, OH), 100 mM NH_4_Cl and 2.5% horse serum in PBS. Then, samples were probed with in-house-made antisera (mouse anti-SAPA, 1:3000 or rabbit antiVSG-221, 1:2000) and/or rat anti-FLAG mAb M2 clone, 1:1000 (Sigma)) diluted in BS. Following extensive washes with PBS, the corresponding Alexa Fluor-conjugated secondary antibodies (Invitrogen, IL) diluted in BS to a final concentration of 2 μg/ml were added. Both primary and secondary antibody incubations were 1h long and all steps were performed at room temperature in a humid chamber. When necessary, DAPI (4’,6-diamidine-2’phenylindole dihydrochloride) was used (CalBiochem, Germany) to visualize the nucleus and kinetoplast before montage with FluorSave reagent (Merk).

For two-color STORM samples, post-labeling fixation was performed as described above in a 6-well polystyrene plate (Corning) and samples were stored in PBS at 4°C until processed. Alexa Fluor 568 and Alexa Fluor 647 were chosen due to the low crosstalk between their emission spectra and their performance for STORM imaging^31^.

### Imaging

The STORM microscope was custom-built around a commercial inverted microscope stand Olympus IX-73 operating in wide-field epifluorescence mode. A 642 nm 1.5 W laser (MPB Communications 2RU-VFL-P-1500–642) and a 532 nm 1.5 W laser (Laser Quantum Ventus 532), both circularly polarized, were used for fluorescence excitation. A 405 nm 50 mW diode laser (RGB Photonics Lambda Mini) was used for reactivating fluorescent molecules. The lasers were focused to the back focal plane of the oil immersion objective Olympus PlanApo 60x NA 1.42. Collected fluorescence light was decoupled from the laser excitation by a quad-edge multiband mirror (Semrock Di03-R405/488/532/635-t1). Further blocking of the illumination lasers was performed with a quad-notch filter NF (Semrock NF03405/488/532/635E-25).

Dual-color simultaneous imaging was performed through emission wavelength discrimination. A dichroic lowpass filter (Chroma ZT647rdc) and two band-pass filters (Semrock 582/75 BrightLine HC and Chroma ET700/75m) were used to separate the fluorescence emission from Alexa Fluor 568 and Alexa Fluor 647, respectively.

Differences in magnification, shear and image rotation between the two channels were corrected prior to acquisition by imaging multicolor beads (Life Technologies Tetraspeck 0.1 μm), performing an affine transformation correction for the overlay of channels^40^. The emission light was expanded with a 2x telescope so that the pixel size of the EMCCD camera (Andor iXon3 897 DU-897D-CS0-#BV) was 133 nm in the sample plane. The camera and lasers were controlled with Tormenta, a custom software developed in the laboratory^41^.

### STORM sample preparation

Coverslips (22 mm) containing fixed parasites were placed in a holder and imaging was performed in STORM imaging buffer (50mM Tris, 10mM NaCl, 10% w/v glucose, pH=8) supplemented with 1 μg/mL glucose oxidase (Sigma-Aldrich), 0.5 ug/mL catalase (SigmaAldrich) and 10 mM MEA (mercaptoethylamine) as oxygen scavenging system.

### Image acquisition

Prior to STORM imaging, conventional fluorescence images of the region of interest were acquired by setting the excitation laser power density to 1-3 W/cm^2^. The channel overlay calibration was performed before each measurement routine. Super-resolution data acquisition was then started by changing the excitation laser power density to 10-20 kW/cm^2^, inducing on-off switching of the fluorophores. Given that the parasites are approximately 1 µm thick, the illumination was set up in epifluorescence mode. Throughout the whole acquisition, the 405 nm laser power was increased whenever the density of single-molecule blinking events decreased below 1 molecule per μm^2^. Generally, it took 20000-25000 frames with an exposure time of 50 ms for each STORM acquisition.

### Image post-processing

Picasso software^42^ was employed for the detection and fitting of single-molecule events using the MLE Integrated Gaussian method^43^. Localizations were filtered based on the global estimated precision (11nm) obtained with the NeNA algorithm^44^. Picasso was also used for rendering super-resolution images. Additionally, redundant cross-correlation (RCC) drift correction was applied to all datasets (in each one, we first corrected the drift via RCC in one of the emission channels to generate a drift-correction file, which was then applied to correct the drift in the second channel to ensure that both channels had the exact same correction). For the subsequent analysis of data, 184 circular areas with variable known diameters were selected from 25 parasites - without overlapping - from the entire set of measured parasites. The selection was unbiased, ensuring that no specific region of the parasite’s structure influenced the choice. Localizations within each selected area were then exported for further processing. The parasite edges were excluded to reduce artifacts caused by 2D axial projections.

### Clustering analysis and simulations

The spatial distributions of mucins and TS were analyzed in a total of 25 parasites derived from three independent replicates. Specifically, the number of parasites analyzed from each replicate was 11, 6, and 8 parasites, respectively. DBSCAN^23^ - Density-Based Spatial Clustering of Applications with Noise - was applied to the lists of STORM localizations obtained. This clustering method was chosen due to the unknown number and shape of the clusters and the presence of isolated localizations in the data (noise). MinPoints parameter was set to a value of 10, following the mean number of switching cycles expected for Alexa Fluor 568 and Alexa Fluor 647^31^. The density-based clustering validation (DBCV)^24^ was employed for hyperparameter tuning, given that we lacked prior information about the features of the domains under study. The DBSCAN search radius (ε) was varied from 10 nm to 40 nm in 1 nm increments, and for each area, the combination of fixed MinPoints and the optimal ε value that yielded the highest validation index was selected.

For the randomization of the experimental clusters, the positions of their centers of mass were randomly relocated inside the selected circular area. This process involved initially loading the cluster data, including positions and mass centers. For each cluster, a random point within the area was generated, and the cluster was then shifted to this new position, ensuring it remained within the circle’s boundary and did not overlap with previously relocated clusters. Overlaps were assessed by computing the convex hulls of the clusters and checking for intersections between them.

In the shuffling method applied to clustered localizations, a mucin dataset from an area was combined with a TS dataset from another area, ensuring that the diameters of the selected areas were the same. This approach was used to study cross metrics, while maintaining the native non-random organization of mucins and TS clusters. To quantify the overlap between clusters and estimate their area of occupancy, we generated an image by rendering each localization with a normalized Gaussian. For each normalized Gaussian, we used a standard deviation of 10 nm (i.e., the localization precision). The pixel size of the rendered images was 5 nm. These images were then thresholded using the standard deviation of the rendered image to create binary masks (σ = 10 nm). The overlap between the masks was determined by counting the number of overlapping pixels. The total occupied areas for the mucin and TS masks were similarly calculated by summing the pixels in each mask. Finally, occupancy percentages were computed as the ratio between occupied areas and the total area.

### Dimensional analysis and fibers simulations

To assess the spatial organization of molecular localizations, we analyzed the functional dependence of the complementary cumulative distribution function (CCDF) of the first neighbor distance (*τ*_1_)^32^. This method exploits the scaling behavior of the number of molecules encountered within a given radius (*τ*), which varies based on the underlying structural organization. For a strictly linear arrangement, the number of encountered molecules scales proportionally to *τ* (n=1), whereas for a random two-dimensional (2D) distribution, it follows *τ*^2^ (n=2). Intermediate cases, such as fibrillary structures with varying degrees of branching or crossing, exhibit scaling exponents in the 1<n<2 range, reflecting deviations from both purely linear and purely random 2D distributions. By quantifying this scaling exponent, we can infer the degree of molecular organization and identify fibrillary arrangements within the dataset.

### Blue Native non-denaturing polyacrylamide gel electrophoresis

Tissue-culture derived *T. cruzi* trypomastigotes were harvested by centrifugation at 3000 g for 10 minutes at 4°C and azido sialylated as described above. Exponentially grown *T. brucei* bloodstream trypomastigotes forms Lister 427 VSG 221 were harvested at 1000 g under the same conditions. The parasites were first washed twice with cold PBS and then resuspended in cold Blue Native lysis buffer (BN-lysis buffer: 20 mM Bis-Tris, 500 mM ε-aminocaproic acid, 20 mM NaCl, 0.2 mM EDTA, pH 8, and 10% glycerol, pH 7), supplemented with 1% (w/v) dodecyl β-D-maltopyranoside (Biosynth) and protease inhibitor cocktail (Sigma). *T. cruzi* trypomastigotes and *T. brucei* bloodstream forms were resuspended to final concentrations of 1.5 × 10⁶ and 1.5 × 10⁵ cells/µL, respectively. Parasite homogenates were kept on ice for 15 minutes and then centrifuged at 10,000 g for 20 min at 4 °C. Native samples were prepared by mixing 4 volumes of the soluble fractions with 1 volume of Non-SDS sample buffer (NSDS-sample buffer: 100 mM Tris-HCl, 150 mM Tris-Base, 0.01875% (w/v) Coomassie G250, 0.00625% (w/v) phenol red, and 10% glycerol, pH 8.5). Equal volumes of each sample were resolved on a NativePAGE™ 3-12% Bis-Tris Gel (Invitrogen), using cathode (15 mM Bis-Tris, 50 mM Tricine, 0.02% (w/v) Coomassie Blue G-250, pH 7.0) and anode buffers (50 mM Bis-Tris, pH 7.0). The electrophoretic run was carried out at 140 V for 45 min at 4 °C. The cathode buffer was then replaced with a light blue cathode buffer (0.1X Coomassie Blue G-250), and the run continued under the same conditions until the dye front reached the bottom of the gel. The resolved proteins were transferred to a PVDF membrane using a transfer buffer (25 mM Tris, 192 mM glycine, pH 8, 20% (v/v) methanol) supplemented with 0.05% (w/v) SDS. Immunoblotting was performed using in-house-made rabbit anti-SAPA (1:3000), rat anti-TolT-A (1:1000) and mouse anti-VSG 221 (1:200) antisera. *T. cruzi* sialylated samples were probed using rat anti-FLAG monoclonal antibody (L5 clone, 1:1000, Biolegend). Secondary antibody probing was performed using goat HRP-conjugated secondary antibodies against rat, mouse, or rabbit immunoglobulins (1:5000, BioLegend). Both primary and secondary antibodies were diluted in blocking buffer Tris-buffered saline (TBS: 50 mM TrisHCl, pH 7.6, and 150 mM NaCl) containing 5% (w/v) non-fat dry milk. Chemiluminescent detection was performed using SuperSignal™ West Pico PLUS reagent (Thermo Scientific™).

## Acknowledgements

F.D.S. and A.M.S. acknowledge the Alexander von Humboldt Foundation. F.D.S thanks Exemys S.R.L. for support with the hardware. J.M. thanks Liliana Sferco and Agustina Chidichimo for help with parasite cultures and Francisco Güaimas for advice on microscopy. We are grateful to Dr. Esteban Erben (IIB-UNSAM, Buenos Aires) for providing *T. brucei* parasite and anti-VSG 221 antibody.

## Author contributions

G.E. performed super-resolution imaging measurements, developed code for the data analysis and analyzed the data and prepared the figures. R.P.P. and M.M.C. prepared the *T. cruzi* samples and performed the biochemical assays. L.F.L performed super-resolution imaging. L.F.L. and A.M.S conceived analysis with G.E.. L.A.M. performed preliminary superresolution measurements. F.D.S, O.E.C and J.M. conceived the project and designed the experiments. G.E., F.D.S., L.F.L and J.M wrote the manuscript with input from all authors. F.D.S and J.M. supervised the project.

## Funding

This work was funded by Consejo Nacional de Investigaciones Científicas y Técnicas (CONICET, Argentina), through grants from the Agencia Nacional de Promoción de la Investigación, el Desarrollo Tecnológico y la Innovación (ANPCyT, Argentina, PICT V 20143729, PICT-2017-0870 and PICT-2021-01216 (FDS) and PICT-2020-00419 (JM)), the Red Federal de Microscopía de Superresolución (FDS), and the National Institute of Allergy and Infectious Diseases (NIAID) of the National Institutes of Health under awards number R01AI104531 (to OC) and R01AI191327 (to JM). The funders had no role in the design of studies, data collection, and analysis, or in the decision to publish or the preparation of the manuscript.

## Data availability statement

All data included in this work along with the open-source code used to analyze the data and generate the figures are publicly available in https://zenodo.org/records/16325107

## Supplemental Material

“Distinct nanoscale membrane organizations of mucins and trans-sialidases in ***Trypanosoma cruzi*”** (Escalante et al.)

**Supplementary Figure 1.**
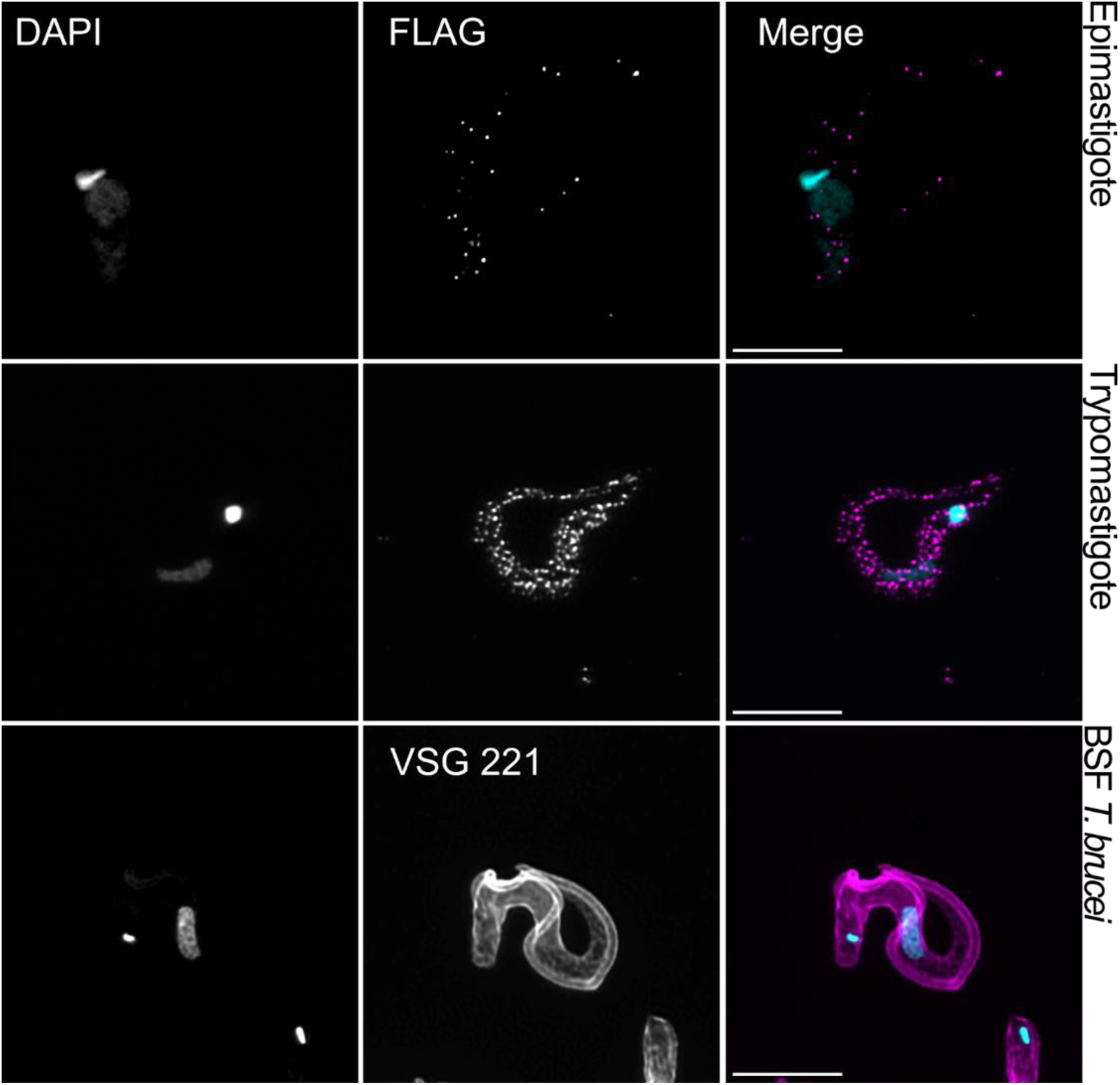
Comparative confocal analysis of surface glycoprotein distribution. Mucin distribution in epimastigote T. cruzi (CL strain, top panels), cell culture-derived trypomastigote T. cruzi (middle panels) and VSG 221 distribution in T. brucei (bloodstream form, bottom panels). All parasites were prepared and PFA-fixed under the same conditions. Images are presented as confocal maximum Z-projections from images acquired with a Zeiss LSM 980 with Airyscan 2. The uniform distribution of VSG serves as a negative control for PFA-induced protein aggregation. Mucins (labeled with the Staudinger-Bertozzi click-chemistry approach and rat monoclonal anti-FLAG) and VSG (immunolabeled with rabbit polyclonal anti-VSG 221) are shown in magenta, while kinetoplasts and nuclei were stained with DAPI and shown in cyan. Scale bars are 5 μm.

**Supplementary Figure 2.**
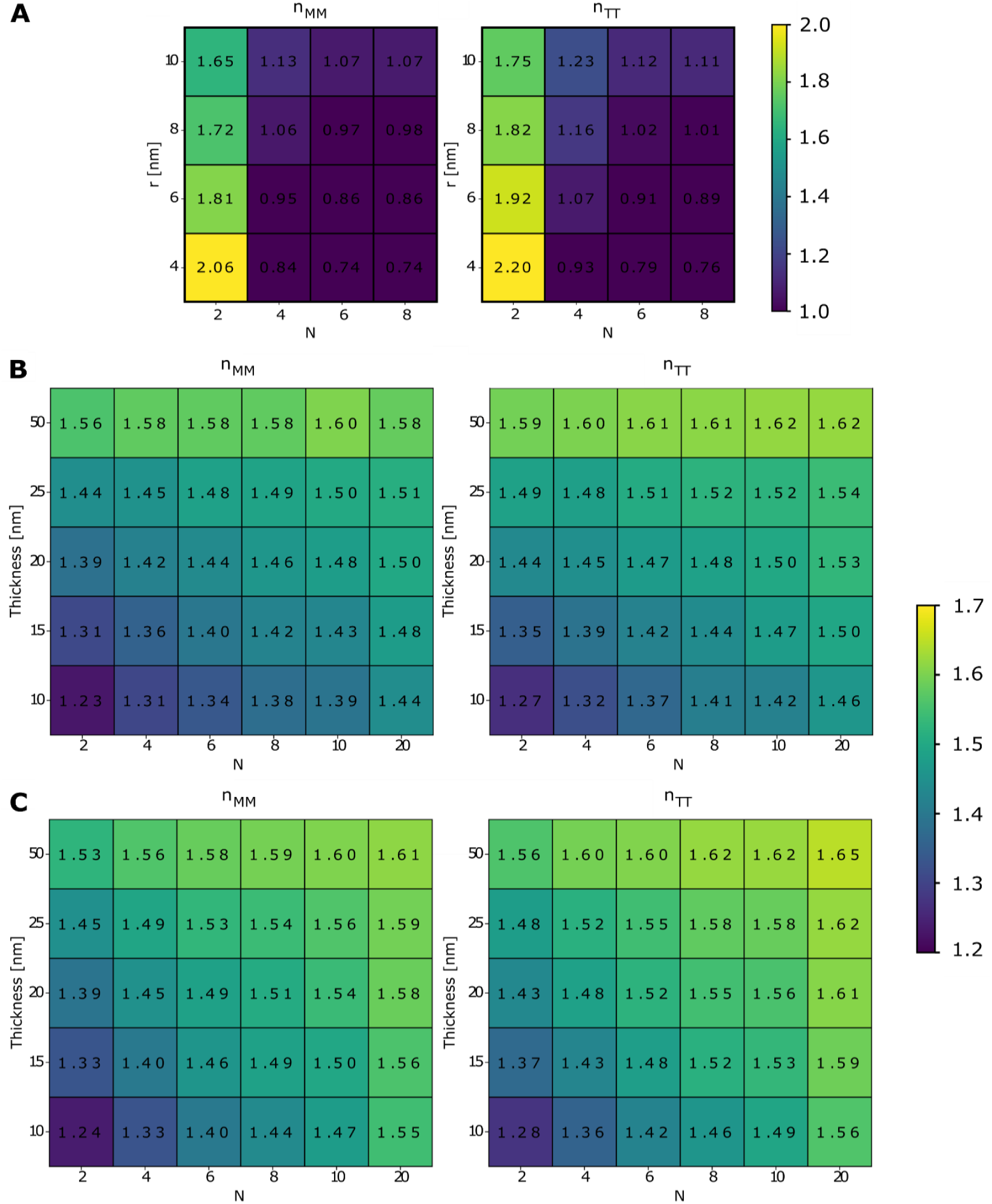
Dimensionality (*n*) analysis for three structural models: **(A)** Circular compartments (r: radius, N: number of circles), evaluating mucin-mucin (n_MM_) and TS-TS (n_TT_) distributions. **(B)** Linear fibers (w: width; N: number of fibers), assessing mucin-mucin (n_MM_) and TS-TS (n_TT_). **(C)** Curved fibers (w: width; N: number of fibers), mucin-mucin (n_MM_) and TS-TS (n_TT_). For each parameter set, 10,000 Monte Carlo simulations were performed across all models.

**Supplementary Figure 3.**
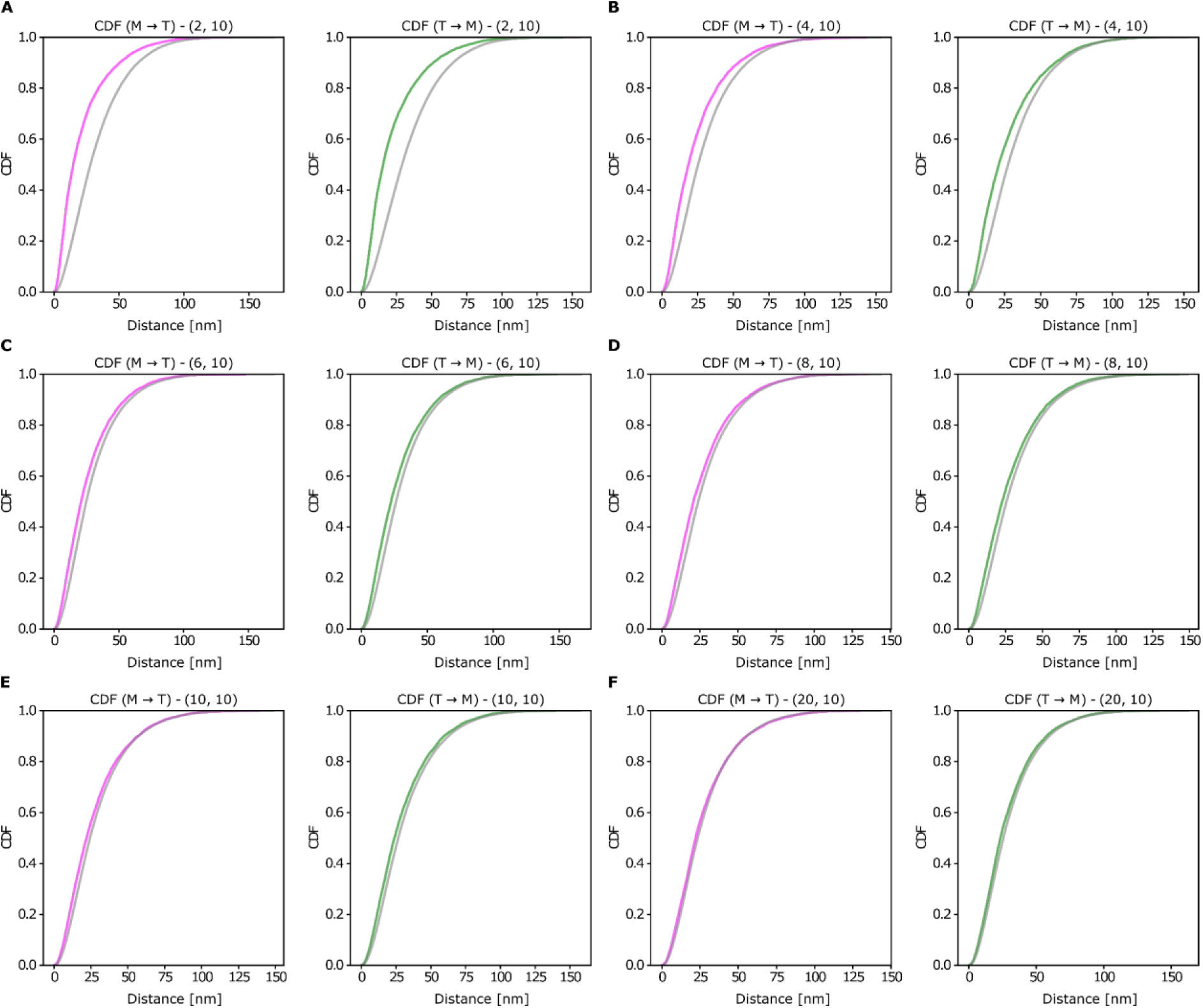
CDFs calculated for the curved fibers model for the pairs (N, thickness = 10) varying the amount N of fibers simulated. Simulated M-T distances are represented in magenta, simulated T-M in green and shuffled M-T and T-M in gray for **(A)** N = 2 curved fibers, **(B)** N = 4 curved fibers, **(C)** N = 6 curved fibers, **(D)** N = 8 curved fibers, **(E)** N = 10 curved fibers and **(F)** N = 20 curved fibers.

**Supplementary Figure 4.**
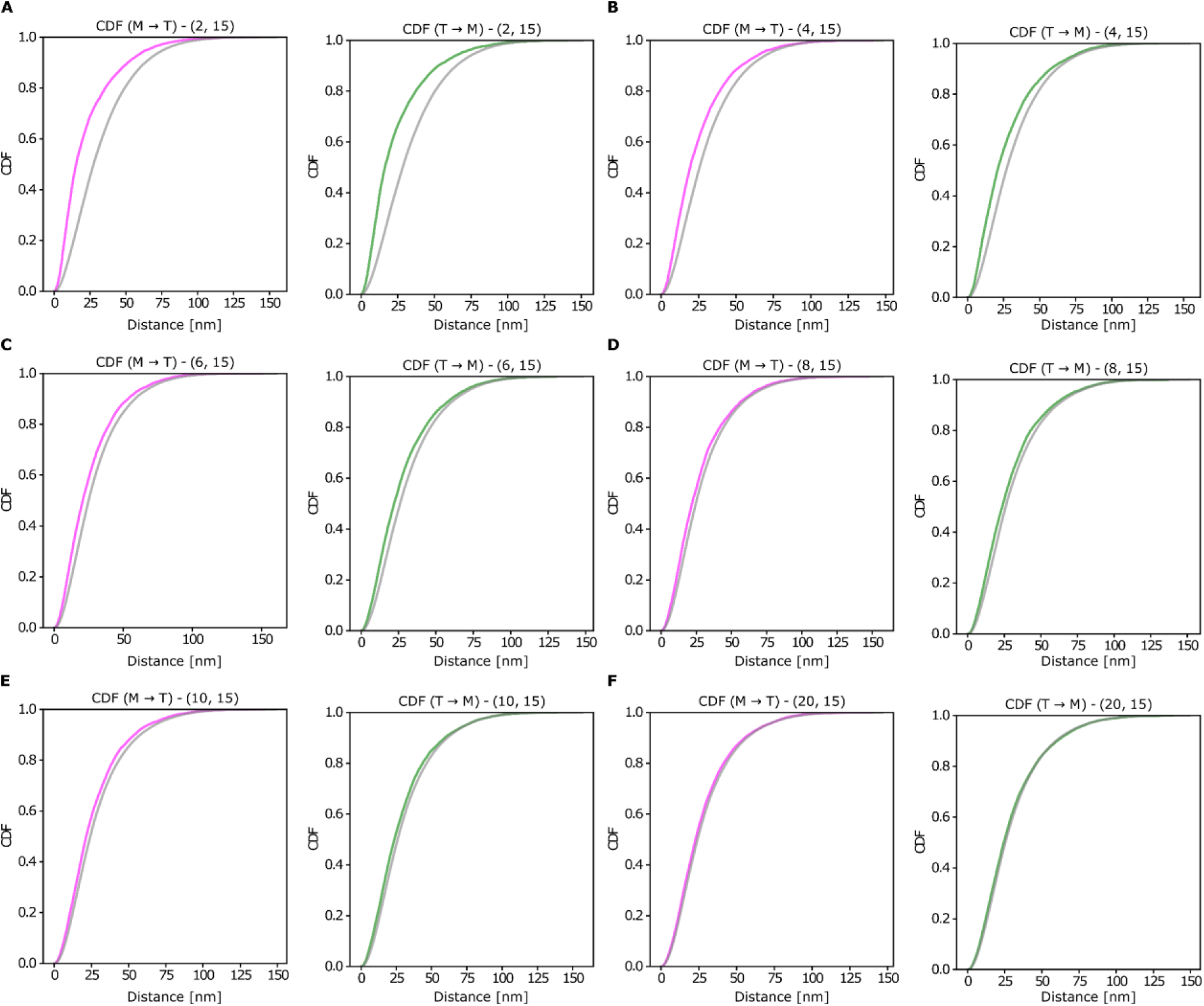
CDFs calculated for the curved fibers model for the pairs (N, thickness = 15) varying the amount N of fibers simulated. Simulated M-T distances are represented in magenta, simulated T-M in green and shuffled M-T and T-M in gray for (A) N = 2 curved fibers, (B) N = 4 curved fibers, (C) N = 6 curved fibers, (D) N = 8 curved fibers, (E) N = 10 curved fibers and (F) N = 20 curved fibers.

**Supplementary Figure 5.**
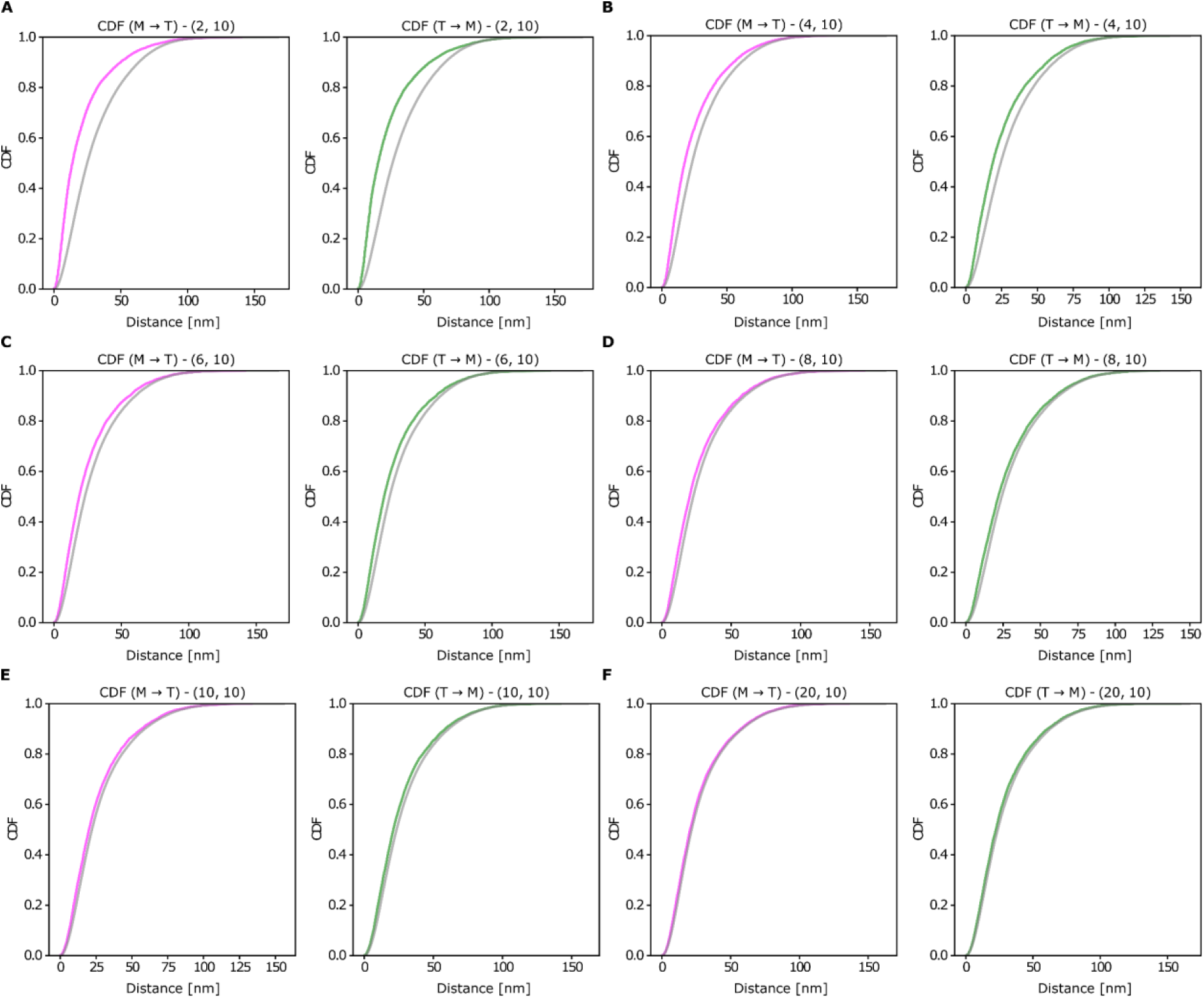
CDFs calculated for the linear fibers model for the pairs (N, thickness = 10) varying the amount N of fibers simulated. Simulated M-T distances are represented in magenta, simulated T-M in green and shuffled M-T and T-M in gray for (A) N = 2 curved fibers, (B) N = 4 curved fibers, (C) N = 6 curved fibers, (D) N = 8 curved fibers, (E) N = 10 curved fibers and (F) N = 20 curved fibers.

**Supplementary Figure 6.**
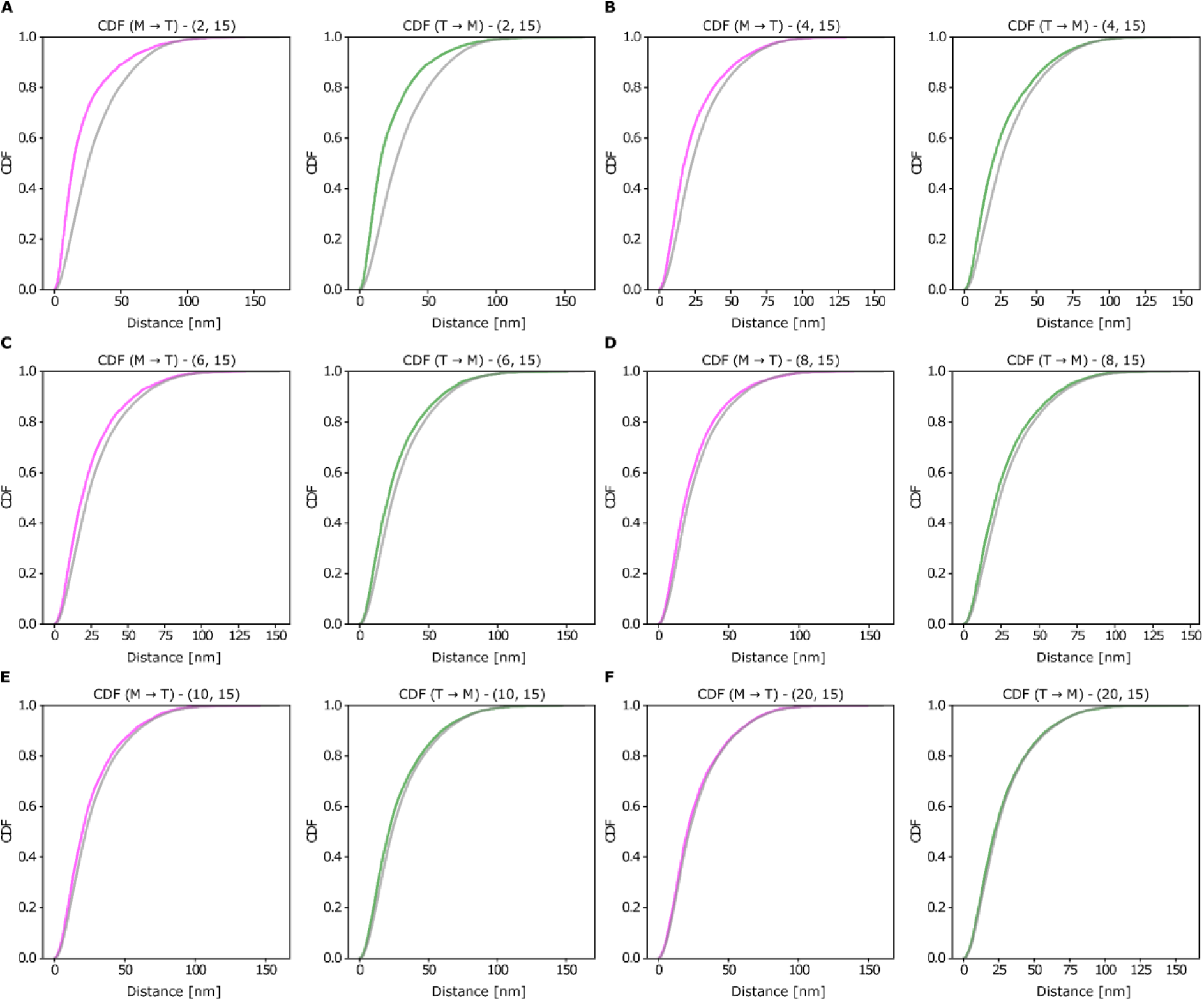
CDFs calculated for the linear fibers model for the pairs (N, thickness = 15) varying the amount N of fibers simulated. Simulated M-T distances are represented in magenta, simulated T-M in green and shuffled M-T and T-M in gray for (A) N = 2 curved fibers, (B) N = 4 curved fibers, (C) N = 6 curved fibers, (D) N = 8 curved fibers, (E) N = 10 curved fibers and (F) N = 20 curved fibers.

## Notes

https://zenodo.org/records/16325107

### Competing Interest Statement

The authors have declared no competing interest.

### Summary of Updates

Improved explanations of the measurements and data analysis. Supplementary Figures included after the main manuscript.

